# Ventral striatal dopamine encodes unique properties of visual stimuli in mice

**DOI:** 10.1101/2022.09.20.508670

**Authors:** L. Sofia Gonzalez, Austen A. Fisher, Shane P. D’Souza, Evelin M. Cotella, Richard A. Lang, J. Elliott Robinson

## Abstract

The mesolimbic dopamine system is an evolutionarily conserved set of brain circuits that plays a role in attention, appetitive behavior, and reward processing. In this circuitry, ascending dopaminergic projections from the ventral midbrain innervate targets throughout the limbic forebrain, such as the ventral striatum/nucleus accumbens (NAc). Dopaminergic signaling in the NAc has been widely studied for its role in behavioral reinforcement, reward prediction error encoding, and motivational salience. Less well characterized is the role of dopaminergic neurotransmission in the response to surprising or alerting sensory events. To address this, we used the genetically encoded dopamine sensor dLight1 and fiber photometry to explore the ability of striatal dopamine release in to encode the properties of salient sensory stimuli in mice, such as threatening looming discs. Here, we report that NAc lateral shell (LNAc) dopamine release encodes the rate and magnitude of environmental luminance changes rather than visual stimulus threat level. This encoding is highly sensitive, as LNAc dopamine could be evoked by light intensities that were imperceptible to human experimenters. We also found that light-evoked dopamine responses are wavelength-dependent at low irradiances, independent of the circadian cycle, robust to previous exposure history, and involve multiple phototransduction pathways. Thus, we have further elaborated the mesolimbic dopamine system’s ability to encode visual information in mice, which is likely relevant to a wide body of scientists employing light sources or optical methods in behavioral research involving rodents.

## INTRODUCTION

The mesolimbic dopamine system is an evolutionarily conserved set of circuits that plays a role in approach and avoidance, appetitive behavior, and reward processing (Wise, 2004; Everitt and Robbins, 2005; Alcantara et al., 2022). In this circuitry, ascending dopaminergic projections from the ventral midbrain, including the ventral tegmental area (VTA), innervate targets throughout the limbic forebrain, such as the ventral striatum/nucleus accumbens (NAc). Dopaminergic signaling in the NAc has been widely studied for its involvement in motivational salience, behavioral reinforcement, and reward prediction error encoding (Schultz et al., 2015; Berridge and Robinson, 2016; Watabe-Uchida et al., 2017; Berke, 2018). Less well characterized is the role of dopaminergic neurotransmission in the response to unpredicted or alerting sensory events, which may encourage investigation or prime motivated behavioral responses to these stimuli (Horvitz, 2000; Bromberg-Martin et al., 2010a; Schultz, 2010). While many previous studies have reported phasic firing of dopaminergic neurons in response to light flashes in laboratory animals (Horvitz et al., 1997; Comoli et al., 2003; Dommett et al., 2005), it is unclear how NAc dopamine release encodes the properties and/or emotional valence of arousing visual stimuli, such as visual threats.

Across a range of species (Ball and Tronick, 1971; Sun and Frost, 1998; Maier et al., 2004; Nakagawa and Hongjian, 2010; Yilmaz and Meister, 2013; Temizer et al., 2015), rapidly approaching objects or looming visual threats elicit automatic defensive or avoidance responses. In mice, presentation of an expanding, overhead, black disc that simulates aerial predator approach (a looming stimulus) promotes rapid escape to an available shelter, followed by long periods of freezing (Yilmaz and Meister, 2013). In our previous work published in eLife (Robinson et al., 2019), mice modeling cognitive dysfunction associated with neurofibromatosis type 1 (NF1) exhibited more vigorous escape in responses to looming stimulus presentation. Additionally, NAc dopamine release evoked by a white light stimulus was enhanced in NF1 model mice, which was correlated with behavioral conditioning abnormalities. Despite the demonstration that white light can induce NAc dopamine release (Robinson et al., 2019; Kutlu et al., 2021), the striatal dopamine response to visual threats is not well characterized in mice. Additionally, it is unknown what visual stimulus characteristics – if any – are encoded by NAc dopamine. Thus, one cannot fully interpret the significance of aberrant responses in neurodevelopmental disease models without a more thorough understanding of visual stimulus encoding by mesolimbic dopamine release in typically developing subjects.

In this Research Article, we sought to probe ventral striatal dopaminergic responses to arousing visual stimuli, including looming visual threats. Given the ability of dopaminergic neurons to signal stimulus saliency (Bromberg-Martin et al., 2010b), we hypothesized that looming discs would induce ‘alerting’ NAc dopamine release whose magnitude would scale proportionately with perceived threat intensity. To test this hypothesis, we utilized the genetically-encoded sensor dLight1 (Patriarchi et al., 2019) to monitor dopamine release in the NAc lateral shell (LNAc) of freely moving adult C57Bl/6J mice with fiber photometry, as performed previously (Robinson et al., 2019). The LNAc was chosen because dopamine release and/or VTA dopaminergic axon terminal activity encode both stimulus valence and prediction errors in this region (de Jong et al., 2018; Robinson et al., 2019; Yuan et al., 2019). Here, we report that LNAc dopamine release reliably reads out unique visual stimulus properties in mice, a phenomenon that is likely relevant to a wide body of scientists employing light sources or optical methods in behavioral research.

## RESULTS

### Dopaminergic responses to looming visual threats

To explore the encoding of visual threats by LNAc dopamine, we first measured dLight1 signals evoked by looming discs (Figure 1A-D) using a custom Bonsai-controlled (Lopes et al., 2015) setup for programmable visual stimulus presentation on an overhead liquid crystal display (LCD) within a light and sound-attenuating chamber (see *Materials and Methods* for details). During photometry recordings, mice were exposed to trains of five overhead, black, looming discs on a light gray background that we empirically determined produce short-latency escape in C57Bl/6J mice (Figure 1E, Video 1), consistent with previous studies (Evans et al., 2018; Yilmaz and Meister, 2013). As controls, we presented mice with trains of black discs that do not reliably evoke defensive responses (Figure 1E), such as a static disc (a fixed 30.5 cm black disc on a light gray background), a receding disc (a black disc that contracted from 30.5 cm to 0 cm on a light gray background), and contrast inverted discs (light gray static, looming, or receding discs on a black background). We observed that looming discs induced low-amplitude dopamine transients at the onset of the first stimulus in each train that – contrary to our hypothesis – were not significantly different from the dLight1 responses to non-threatening static and receding discs (Figure 1B-C). Surprisingly, repeating these experiments with contrast-inverted discs that do not induce escape (Figure 1E) evoked ~3 to 6-fold greater dopamine release than black discs (Figure 1B-C). This raised the possibility that LNAc dopamine release tracks stimulus brightness rather than threat intensity.

**Figure 1.**
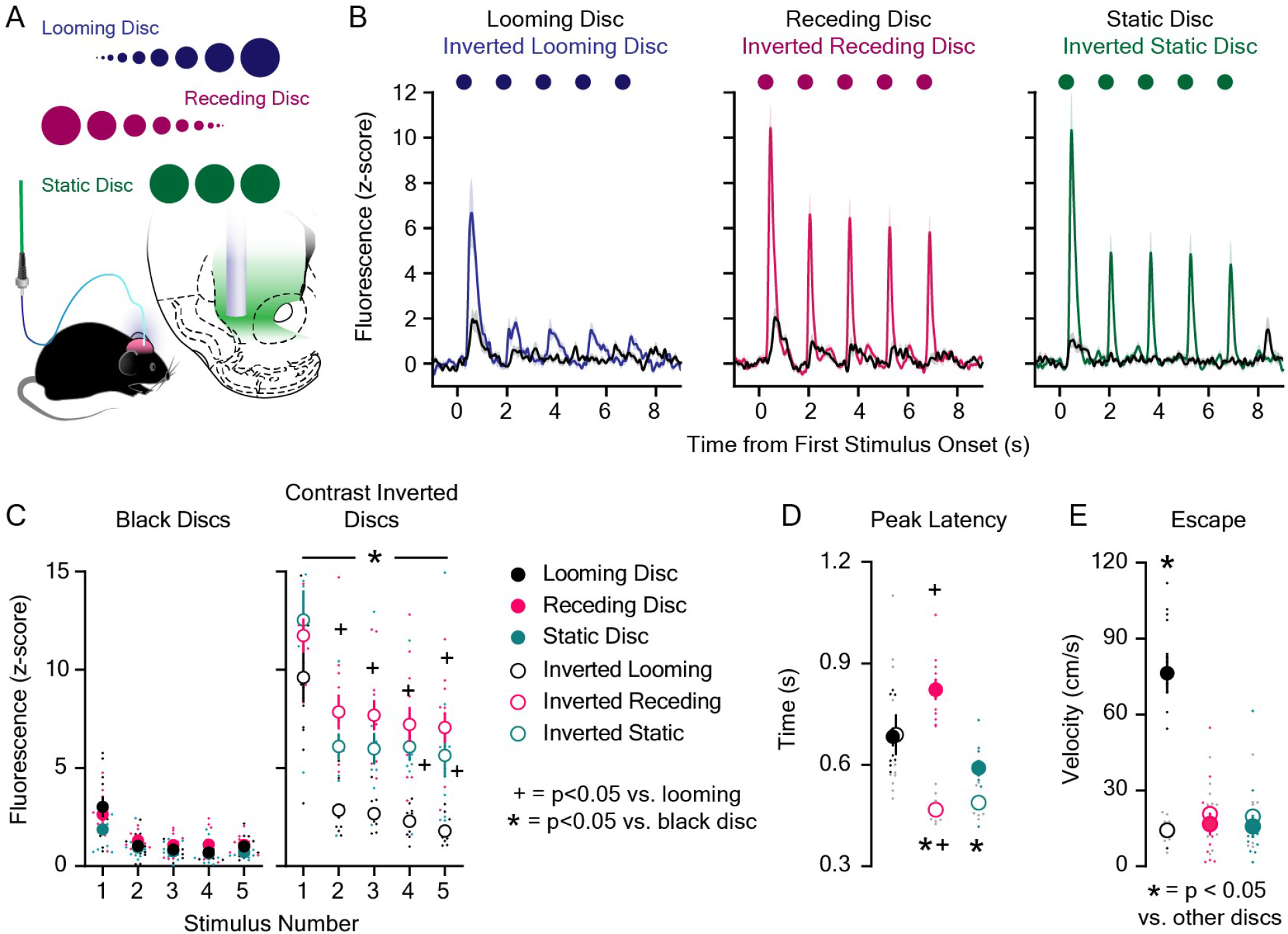
LNAc dopaminergic encoding of visual threats. **A.** Fluorescent dopamine signals were recorded with dLight1 and fiber photometry in the nucleus accumbens lateral shell (LNAc) during presentation of looming and control discs. **B.** Average dLight1 response to trains of five black or contrast inverted discs ± standard error of the mean (SEM). **C.** The dLight1 response to black or inverted discs was dependent on disc color/background, disc type (static vs. looming vs. receding), and stimulus number (n = 11; 3-way repeated measures ANOVA; F_8,240_ = 2.02, p_disc background x disc type x stimulus number_ = 0.045; F_1,240_ = 143.92, p_background_ < 0.001; F_2,240_ = 150.32, p_disc type_ < 0.001; F_4,240_ = 192.64, p_stimulus number_ < 0.001). Bonferroni *post hoc* tests revealed that contrast inverted discs evoked more dopamine than black discs. Contrast inverted looming discs evoked less dopamine than inverted static and receding discs after the first presentation. **D.** dLight1 transient peak latency was dependent on disc color/background and disc type (n = 11; 2-way repeated measures ANOVA; F_2,20_ = 64.78, p_disc background x disc type_ < 0.001; F_1,20_ = 25.69, p_background_ < 0.001; F_2,20_ = 7.58, p_disc type_ = 0.01). Bonferroni *post hoc* tests showed that contrast inverted static and receding discs evoked transients with shorter latency compared to black discs. Additionally, transients evoked by contrast inverted receding discs had shorter latency than contrast inverted looming discs. E. Escape velocity following overhead disc presentation was dependent on disc color/background and disc type (n = 12; 2-way repeated measures ANOVA; F_2,22_ = 49.28, p_disc background x disc type_ < 0.001; F_1,22_ = 18.38, p_background_ = 0.001; F_2,22_ = 28.89, p_disc type_ < 0.001). Bonferroni *post hoc* tests showed that black looming discs induced greater escape velocity than all other overhead discs. For panels C and D, * indicates p < 0.05 vs. black disc of the same type (e.g. black static disc vs. contrast inverted static disc); + indicates p < 0.05 vs looming disc of the same color (e.g. black looming vs. black receding disc). For panel E, * indicates p < 0.05 vs. other overhead discs.

### Dopaminergic responses to rapid changes in environmental lighting conditions

Because inverted looming discs, in which the number of bright overhead pixels ramps as the disc expands, produced lower amplitude (Figure 1C), longer latency (Figure 1D) dLight1 responses than static or receding inverted discs with an instantaneous pixel change, we hypothesized that LNAc dopamine may encode the rate of change of dark-to-light transitions. To test this possibility, we exposed mice to full screen, instantaneous transitions from black to light gray during dLight1 recordings, which eliminated disc edge motion as a contributing visual stimulus property. We found that instantaneous dark-to-light transitions produced a high amplitude (10.38 ± 0.43 z-score), short duration (full width a half maximal amplitude: 143 ± 9.7 ms) dopamine transient that peaked 434 ± 3.3 ms after transition onset (Figure 2A). Lengthening the dark-to-light transition time (i.e. the fade-in time) to full screen illumination (Figure 2B) non-linearly decreased the magnitude of the dLight1 peak and increased the peak latency (Figure 2C). For transition times less than ~500 ms, the dopamine peak latency closely matched the fade-in time, above which peak response occurred hundreds of milliseconds to seconds before full field illumination was reached (Figure 2C). When transition times were greater than 1 s, evoked dLight1 transients were often too small to accurately resolve from the fluorescent baseline for individual mice. However, averaging the fluorescence trace from all mice prior to peak detection allowed signals to be resolved for longer transition times. Thus, results are presented as both the fluorescence peak(s) derived from the photometry trace averaged across all mice (Figure 2C) and individual mice (Figure 2-Figure Supplement 1), which showed high concordance for transition times of 1 second or less (Figure 2-Figure Supplement 1). No dLight1 response was reliably evoked by a ten-second dark-to-light transition despite the stimulus ramping to the same number of bright pixels as trials with shorter transition times (Figure 2B).

**Figure 2.**
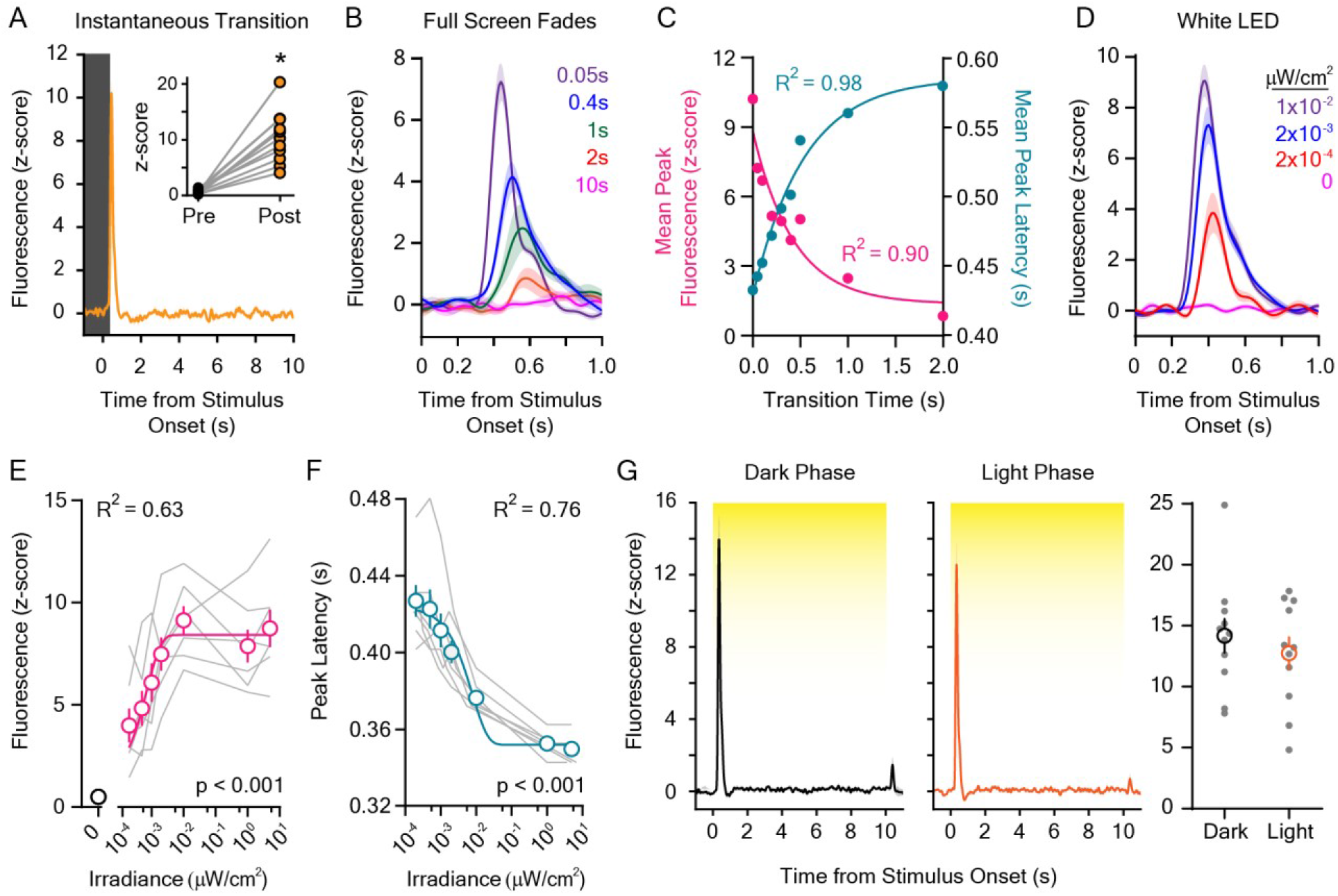
Dopaminergic responses to rapid dark to light transitions. **A.** Instantaneous LCD screen transitions from dark to light evoked rapid dopamine release at stimulus onset when compared to the pre-stimulus baseline (inset: baseline and stimulus-induced dLight1 peak values for individual mice; n = 11; paired t-test; t_10_ = 7.01, p < 0.001). **B.** dLight1 responses to the onset of LCD screen dark to light transitions at different transition lengths (0.05 – 2.0 s) ± SEM. **C.** The magnitude (*pink*) and latency (*teal*) of dopaminergic responses to dark-to-light transitions varied non-linearly depending on transition speed (peak amplitude: one-phase exponential decay, y_0_ = 8.84 z-score, plateau = 1.36 z-score, tau = 0.43 s, R^2^ = 0.90; peak latency: one-phase exponential association, y_0_ = 0.43 ms, plateau = 0.59 ms, tau = 0.54 s, R^2^ = 0.98). **D.** dLight1 responses to the onset ten-second white LED stimuli across a range of irradiances (0 μW/cm^2^ – 0.01 μW/cm^2^) ± SEM. E. The magnitude of the dopaminergic response to 10s white LED stimuli was dependent on the stimulus irradiance (n = 7; 1-way repeated measures ANOVA; F_7,42_ = 38.79, p < 0.001). Data shown with a one-phase exponential association fit (y_0_ = 1.20 z-score, plateau = 0.84 z-score, tau = 0.00074 μW/cm^2^, R^2^ = 0.63). **F.** The latency of the dopaminergic response to 10 s white LED stimuli was dependent on the stimulus irradiance (n = 7; 1-way repeated measures ANOVA; F_6,36_ = 47.35, p < 0.001). Data shown with a one-phase exponential decay fit (y_0_ = 0.42 ms, plateau = 0.35 ms, tau = 0.0079 μW/cm^2^, R^2^ = 0.76). G. The dopaminergic response to 5.0 μW/cm^2^ white light was not different (*right*) if measured at the beginning of the vivarium dark (*left*) or light (*center*) phase of the day-night cycle (n = 11; paired t-test; t_10_ = 1.27, p = 0.23). In all panels, * indicates p < 0.05.

Next, we examined whether LNAc dopamine release also reads out the magnitude of environmental lighting changes by measuring the dLight1 response to ten-second, instantaneous exposures to white light across a range of intensities (0.2 nW/cm^2^ – 5.0 μW/cm^2^, measured at mouse level) generated by a light emitting diode (LED) (Figure 2D-F). High irradiance LED illumination (5 μW/cm^2^) evoked a dLight1 transient at stimulus onset (Figure 2D) that was similar to transients evoked by the LCD monitor (Figure 2A-B; irradiance at mouse level: 11 μW/cm^2^). This response did not depend on the time of testing within the vivarium day-night cycle (Figure 2G) and was larger than the response to auditory tones (80 dB; 1 – 16 kHz; Figure 2-Figure Supplement 1). When LED irradiance was reduced, we observed an intensity-dependent decrease in the magnitude of the dLight1 peak and increase in the response latency (Figure 2E-F), consistent with Bloch’s law of temporal summation in mammalian photoreceptors (Scharnowski et al., 2007; Donner, 2021). Significant dopaminergic responses were observed at all irradiances tested, including 0.2 nW/cm^2^, which was not perceptible to the human experimenter. As a point of reference, the lock screen of a Samsung S21 smart phone on the lowest brightness setting had an irradiance of 20 nW/cm^2^ when placed in the same position as the white LED. Likewise, time-locked dopamine release could be evoked by simply uncovering the enclosure peephole that allows users to observe mouse behavior (irradiance: 17 nW/cm^2^; Figure 2-Figure Supplement 1). These results were not likely caused by mouse movement, as illumination of a white LED that was 1000-fold more intense (5 mW/cm^2^) than the highest irradiance tested had little effect on behavior when freely exploring mice entered a target zone within a dark arena (Video 2). Thus, LNAc dopamine release is sensitively evoked by ambient light and reliably encodes the speed of these lighting transitions over short timescales.

### Dopaminergic responses to repeated light stimuli

Previous literature suggests that dopaminergic neuron firing in response to sensory events habituates as the novel stimulus becomes familiar (Schultz, 1998). In order to test if the dLight1 response to white light is affected by a history of previous exposures, we exposed mice to twenty consecutive one-second white light pulses over five trials (100 pulses total) across a range of interstimulus intervals (ISIs; 10 ms to 10 s; Figure 3A). We found that light-evoked dopamine transient magnitude decayed logarithmically as a function of the ISI duration (Figure 3B). When the ISI was short (e.g. 10 – 100 ms), dLight1 responses habituated rapidly. This is exemplified by the dopaminergic response to 40 Hz light flicker, which is used therapeutically to enhance neural activity in the context of Alzheimer’s disease (Singer et al., 2018). Presentation of a sixty-second 40 Hz white LED flicker (5.0 μW/cm^2^ irradiance, 50% duty cycle) induced a dopamine transient only at stimulus onset (Figure 3C) that was indistinguishable from the response to constant illumination (Figure 2D). Conversely, in shorter experiments when the ISI was long (100 s), no habituation of the dopamine response to light was observed from stimulus-to-stimulus (Figure 3D). Repeating this experiment immediately after three hundred consecutive one-second exposures with a one-second ISI produced an ~20% reduction in the dLight1 response to 100 s ISI light; however, this effect did not reach statistical significance (p = 0.08) (Figure 3D). Therefore, habituation of the dopamine response to repeated light stimuli is much more strongly influenced by stimulus frequency than the total number of previous exposures.

**Figure 3.**
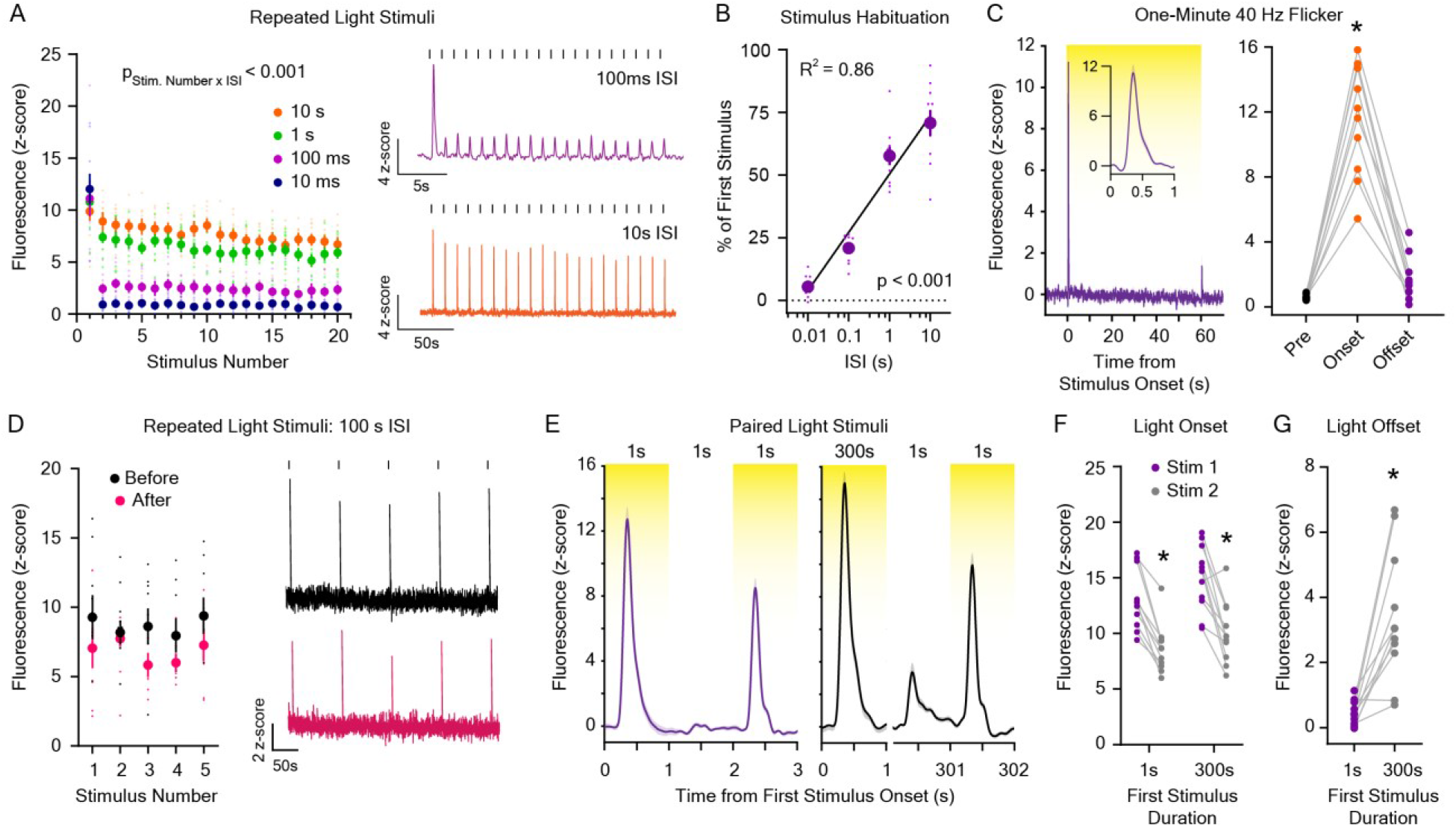
Dopaminergic responses to repeated light stimuli. **A.** (*Left*) Dopamine release evoked by 20 one-second white LED stimuli was reduced with repeated exposures and was dependent on the interstimulus interval (ISI; 10 ms – 10 s; n = 9; 2-way repeated measures ANOVA; F_57,456_ = 9.54, p_stimulus number x interstimulus interval_ < 0.001; F_19,456_ = 72.98, p_stimulus number_ < 0.001; F_3,456_ = 63.91, p_interstimulus interval_ < 0.001). (*Right*) Averaged dLight1 fluorescent traces showing the dopaminergic response to 20 one-second white LED light pulses with a 100 ms ISI (*purple*) or 10 s ISI (*orange*). **B.** Total habituation of the peak dLight1 response to repeated stimuli (shown as the peak response to the 20^th^ stimulus as a percentage of the 1^st^ stimulus) is dependent on the duration of the interstimulus interval (n = 9; 1-way repeated measures ANOVA; F_3,24_ = 104.0, p < 0.001). Data shown with a semi-log fit (y-intercept: 48.92%, slope: 22.58 %s^-1^, R^2^ = 0.86). **C.** (*Left*) Averaged dLight1 trace showing LNAc dopamine evoked by a 60 second presentation of 40 Hz white LED flicker (*inset:* response during the first second after stimulus onset) ± SEM. (*Right*) 40 Hz flicker only evoked significant dopamine release at stimulus onset (n = 10; 1-way repeated measures ANOVA; F_2,18_ = 100.4, p < 0.001). Bonferroni *post hoc* tests confirmed that the dLight1 peak at LED onset was greater than baseline and offset responses, which did not differ from each other (p = 0.09). **D.** (*Left*) No habituation in the peak dLight1 response to repeated 1 s white LED stimuli was observed when the ISI was sufficiently long (100 s) – both before and after presentation of 300 one-second LED stimuli with a one-second ISI (n = 7; 2-way repeated measures ANOVA; F_4,24_ = 0.73, p_stimulus number x exposure history_ = 0.52; F_4,24_ = 1.01, p_stimulus number_ = 0.40; F_1,24_ = 4.61, pexposure history = 0.08). (*Right*) Averaged dLight1 fluorescent traces showing the dopaminergic response to 5 one-second white LED light pulses with a 100 s ISI before (*black*) or after (*pink*) 300 one-second LED stimuli. E. Averaged dLight1 fluorescent traces showing the dopaminergic response to a 1 s white LED stimulus one second after a 1 s (*left*) or 300 s (*right*) preconditioning stimulus ± SEM. **F.** The dLight1 response to a 1 s white LED test stimulus was not dependent on the length of the preconditioning stimulus (n = 11; 2-way repeated measures ANOVA; F_1,10_ = 0.27, p_initial stimulus length x stimulus Number_ = 0.61; F_1,10_ = 3.83, pinitial stimulus length = 0.08; F_1,10_ = 55.10, p_stimulus number_ < 0.001). Bonferroni *post hoc* tests revealed that the dLight1 response to the test stimulus onset was significantly smaller than the response to the onset of the preconditioning stimulus, regardless of its duration. There was no difference between the dLight1 response to the onset of the preconditioning (p = 0.11) or test stimulus (p = 0.40) between experiments. G. The dLight1 response to light offset was larger for a 300 s light stimulus compared to a 1 s light stimulus (n = 11; paired t-test; t_10_ = 4.91, p < 0.001). In all panels, * indicates p < 0.05.

During repeated stimulus experiments, the greatest reduction in the peak LNAc dopamine response to light stimuli occurred between the first and second light pulse in each stimulus train. In order to further characterize this phenomenon, we varied the duration of the first stimulus to determine if the total amount of initial light exposure modulates the dopaminergic response to a subsequent stimulus (Figure 3E). We found that the dopaminergic response to a 1 second white LED test stimulus was not significantly different when preceded by either a 300 second or 1 second preconditioning light stimulus one second earlier (Figure 3F). No difference in dLight1 response to the preconditioning stimulus was observed between conditions (Figure 3F). We did observe, however, that the 300-second preconditioning stimulus produced a dopaminergic response at light offset, whereas the 1-second preconditioning stimulus did not (Figure 3G). Taken together, our findings indicate that ISI is a more significant determinant of stimulus-to-stimulus dopamine release habituation than light stimulus duration.

### Wavelength and photoreceptor contributions to the dopaminergic response to light

In these and previous experiments (Robinson et al., 2019), we employed a white LED light to induce striatal dopamine release; however, this light source is composed of multiple wavelengths throughout the visible spectrum. Therefore, we next investigated if light-evoked dopamine release exhibits wavelength specificity. This is additionally germane given the widespread use of molecular and optical technologies in rodents that require delivery of specific wavelengths of visible light in order to probe neural activity, structure, or biology (Fenno et al., 2011; Resendez and Stuber, 2015; Sabatini and Tian, 2020). In order to determine if the dLight1 response varied by wavelength, we measured dopamine release induced by ten-second exposures to environmental ultraviolet (UV; 360 nm), blue (475 nm), green (555 nm), red (635 nm), and far-red (730 nm) light across a 100,000-fold range of irradiances (1 nW/cm^2^ to 100 μW/cm^2^). These experiments revealed broad sensitivity of the mesolimbic dopamine system to light across the visual spectrum (Figure 4A-B). The dopamine response was least sensitive to UV and red light when the irradiance was low (1 nW/cm^2^; Figure 4A-B), and far-red light (730 nm) only induced dopamine release when the irradiance was high (100 μW/cm^2^; Figure 4-Figure Supplement 1). The ability of red light to induce dopamine release at intensities as low as 0.1 μW/cm^2^ is consistent with research that rodents are better at perceiving red wavelengths than is commonly acknowledged (Danskin et al., 2015; Nikbakht and Diamond, 2021; Vinberg et al., 2019). Whereas the dLight1 response to UV and red light was irradiance-dependent, the response to blue and green light remained robust across the entire irradiance range (Figure 4A). These experiments indicate that the mesolimbic dopamine system is responsive to all visible wavelengths yet is most sensitive to blue and green light.

**Figure 4.**
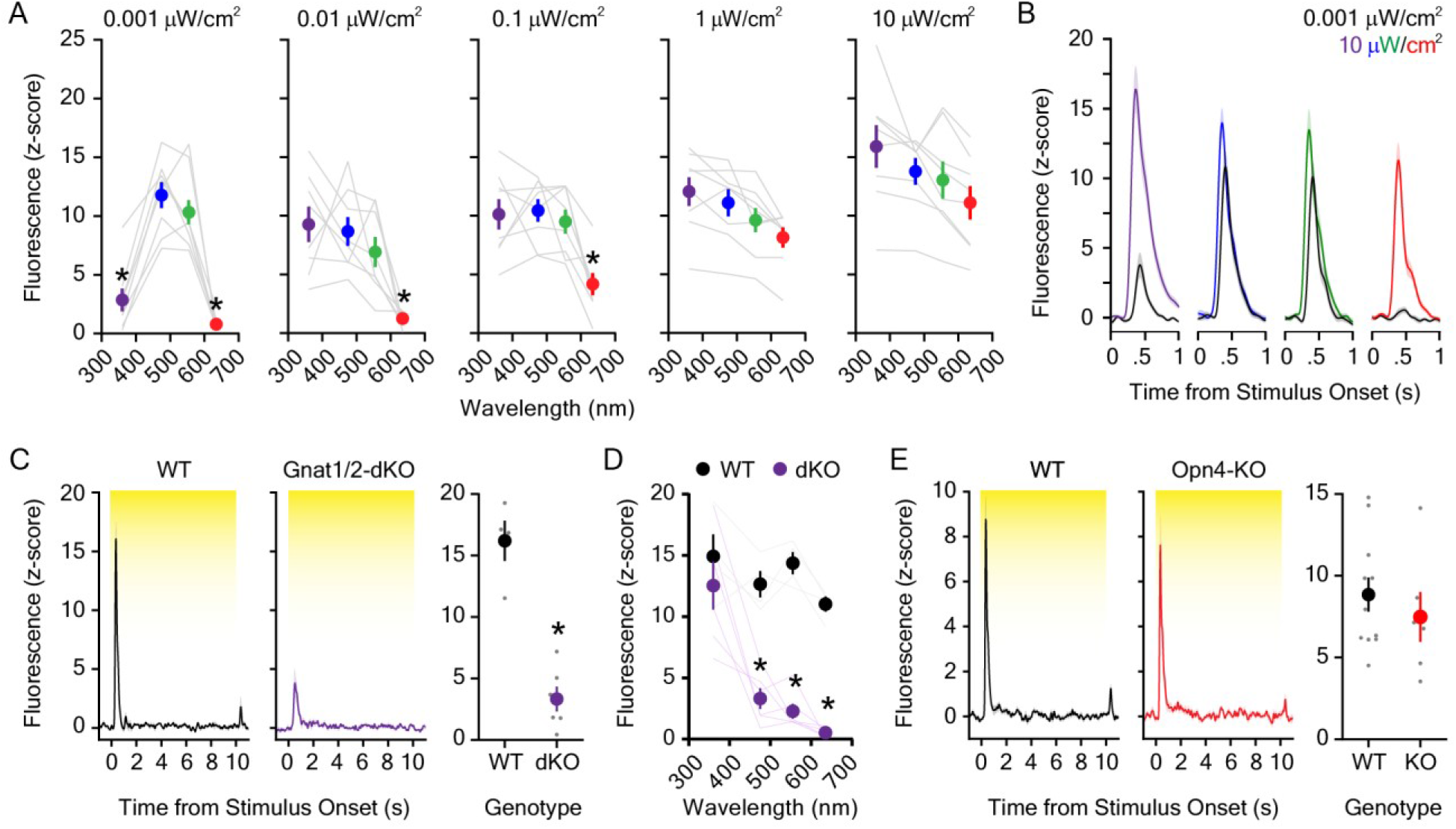
Dopaminergic responses to individual wavelengths across the visual spectrum. **A.** The dopaminergic response to UV (360 nm), blue (475 nm), green (555 nm), and red (635 nm) LED light was wavelength and irradiance-dependent (n = 8; 2-way repeated measures ANOVA; F_12,84_ = 9.63, p_wavelength_ x irradiance < 0.001; F_3,84_ = 37.59, p_wavelength_ < 0.001; F_4,84_ = 10.08, p_irradiance_ = 0.004). Bonferroni *post hoc* tests revealed that dopamine evoked by UV and red light was smaller than blue and green wavelengths at the lowest irradiance tested (0.001 μW/cm^2^). The dLight1 response to the red LED was also significantly lower than blue and green LEDs at irradiances of 0.01 μW/cm^2^ and 0.1 μW/cm^2^. For comprehensive reporting of all significant *post hoc* tests across irradiances and wavelengths, see the provided Supplemental Data and Statistical Analysis file. **B.** Averaged dLight1 trace showing LNAc dopamine evoked by either 0.001 μW/cm^2^ (1 nW/cm^2^) or 10 μW/cm^2^ UV, blue, green, or red LEDs ± SEM. **C.** The dopaminergic response to 5.0 μW/cm^2^ white light was significantly reduced in *Gnat1/2* double knockout (dKO) mice relative to wildtype controls (n_WT_ = 4, n_KO_ = 6; unpaired t-test; t_8_ = 7.08, p < 0.001). **D.** The reduction in the dLight1 response to 10 μW/cm^2^ light in Gnat1/2 dKO was wavelength dependent (2-way repeated measures ANOVA; F_3,24_ = 7.02, p_genotype x wavelength_ = 0.002; F_3,24_ = 17.54, p_wavelength_ = 0.003; F_1,24_ = 85.80, p_genotype_ < 0.001). Bonferroni *post hoc* tests revealed that the dLight1 response to blue (475 nm), green (555 nm), and red (635 nm) light was lower in Gnat1/2 mice relative to wildtype littermates. E. The dopaminergic response to 5.0 μW/cm^2^ white light was not different in *Opn4* (melanopsin) knockout mice relative to wildtype controls (n_WT_ = 11, n_KO_ = 6; unpaired t-test; t_15_ = 0.75, p = 0.46).

The mouse visual system utilizes numerous opsin proteins for image-forming and non-imaging forming phototransduction with unique wavelength sensitivities. These include the rod opsin rhodopsin for scoptopic vision (λ_max_ ~500 nm) and short (λ_max_ ~360 nm) and medium/long wavelength (λ_max_ ~508 nm) cone opsins for photopic vision. Additionally, melanopsin (λ_max_ ~480 nm) is expressed in intrinsically photosensitive retinal ganglion cells (ipRGCs) that mediate circadian entrainment, the pupillary light reflex, and light-regulated changes in mood (Panda et al., 2003; Hattar et al., 2003; Fernandez et al., 2018). While it has been hypothesized that ipRGCs engage VTA dopamine neurons via hypothalamic intermediates (Zhang et al., 2021), the role of melanopsin in the dopaminergic response to light is unknown. In order to parse the role of visual opsins versus melanopsin in the mesolimbic response to dark-to-light transitions, we performed LNAc dLight1 recordings in *Opn4* (melanopsin) knockout mice and *Gnat1/Gnat2* double knockout mice (*Gnat1/2-dKO*). *Gnat1 and 2* knockout mice lack expression of rod and cone α-transducin, respectively, and exhibit loss of signal transduction through these photoreceptors (Deng et al., 2009; Yao et al., 2018). Compared with wildtype littermates, *Gnat1/2-dKO* mice displayed a robust reduction in the dopaminergic response to a 5 μW/cm^2^ white LED (Figure 4C) and an increase in the dLight1 response latency (Figure 4-Figure Supplement 1). Light-evoked dopamine release was not abolished, however, in these mice (Figure 4C). Spectral analysis indicated that *Gnat1/2-* dKO mice retain sensitivity to UV light (Figure 4D, Figure 4-Figure Supplement 1), which may be indicative of residual cone-based vision (Allen et al., 2010). Conversely, loss of melanopsin expression in *Opn4* knockout mice (Panda et al., 2002) did not affect the dLight1 response to the white light stimulus (Figure 4E, Figure 4-Figure Supplement 1). These findings indicate that light-evoked dopamine release is rod and cone-dependent and does not involve melanopsin.

## DISCUSSION

In these investigations, we used the genetically-encoded dopamine sensor dLight1 to demonstrate that LNAc dopamine release can encode rapid changes in luminance but not looming threat intensity. We found that rapid dark-to-light transitions evoked time-locked dopamine responses at stimulus onset at irradiances as low as 0.2 nW/cm^2^, which is in line with findings that mice see over a 100 million-fold range of light intensity beginning at ~4 μcd/m^2^ (Umino et al., 2008). The magnitude of these dopaminergic responses was highly dependent on light stimulus frequency and transition rate rather than duration or novelty. In fact, high amplitude LNAc dLight1 responses to a white LED persisted after hundreds of exposures and months of regular testing. Although mesolimbic dopamine systems regulate wakefulness (Eban-Rothschild et al., 2016) and exhibit circadian oscillation (Korshunov et al., 2017), the time of testing did not appear to be a significant contributor to our findings. Sudden dark-to-light transitions are highly salient to nocturnal rodents that must avoid detection by visual predators (Thompson et al., 2010), so it is possible that the dopaminergic response to light represents a specialized saliency signal. Previously, phasic dopamine responses to unexpected, unconditioned sensory events have been conceptualized in the context of novel action discovery (Redgrave and Gurney, 2006), the need to alert to stimuli that require motivated responses (Schultz and Romo, 1990; Horvitz, 2000), mechanisms of associative learning (Lisman and Grace, 2005), etc. Thus, further studies will be needed to firmly establish the ethological and neurobiological importance of dopaminergic responses to environmental light.

We also found that LNAc dopamine is broadly evoked by wavelengths across the visual spectrum. Given the high proportion of rods in the mouse retina (~97% of photoreceptors) (Jeon et al., 1998) and the reduced sensitivity of dopaminergic responses to 360 and 635 nm light at lower irradiances, it is probable that rod-based phototransduction is primarily responsible for visually-evoked dopamine release under dim (scotopic) lighting conditions. Conversely, rod and cone opsins likely contributed to dLight1 signals in the photopic range. These hypotheses are supported by our observation that genetic disruption of rod and cone-based signaling in *Gnat1/2-* dKO mice substantially attenuated the dopaminergic response to light. *Gnat1/2-dKO* mice retained sensitivity to high irradiance UV light, which was most likely caused by incomplete loss of cone-based vision in this model (Allen et al., 2010). We cannot, however, rule out the involvement of UV-sensitive non-visual opsins in our observed findings, such as neuropsin (*Opn5*), which is maximally activated by 380 nm light (Tarttelin et al., 2003). Neuropsin-expressing retinal ganglion cells project to multiple limbic regions (Sasaki et al., 2021), and this opsin promotes thermogenesis via intrinsically light-sensitive glutamatergic neurons in the preoptic area (Zhang et al., 2020). While melanopsin-expressing ipRGCs are hypothesized to engage VTA outputs via a disynaptic circuit involving the preoptic area (Zhang et al., 2021), we found that *Opn4* knockout had no effect on the ability of light to evoke LNAc dopamine. Given that ipRGCs receive rod and cone input via the retinal synaptic network (Güler et al., 2008; Lall et al., 2010; Altimus et al., 2010), it is possible that these neurons contribute to light-evoked dopamine release independent of melanopsin. Thus, functional lesioning studies will be required to elucidate the role of non-image forming visual pathways in the dopaminergic encoding of visual stimuli.

Visual information is conveyed from the retina to the brain via the axons of retinal ganglion cells that synapse in downstream nuclei to mediate image processing, circadian entrainment, pupillary reflexes, gaze orientation, etc. (Peirson et al., 2018). While thalamocortical visual pathways are required for conscious visual perception, neither the primary visual cortex (V1) nor the visual thalamus (e.g. lateral geniculate nucleus) significantly innervate ventral midbrain dopamine neurons (Watabe-Uchida et al., 2012). Previous work by Redgrave and colleagues suggest that dopaminergic responses to light are driven by the superior colliculus (SC) (Comoli et al., 2003; Dommett et al., 2005; Takakuwa et al., 2017), which receives direct input from retinal ganglion cells (Dhande and Huberman, 2014) in its superficial layers and promotes motivated behavior via deep motor-output layers (Branco and Redgrave, 2020). SC glutamatergic projection neurons directly synapse onto VTA (Solié et al., 2022) and substantia nigra pars compacta dopamine neurons (Huang et al., 2021), both of which project to the LNAc (Beier et al., 2015; Poulin et al., 2018). Likewise, optogenetic stimulation of SC neuron somata is sufficient to evoke LNAc dopamine release *in vivo* (Robinson et al., 2019). While these observations support a role for the SC in dopaminergic responses to light, the relative contribution of different visual processing centers to our findings is an important area of future study.

Mesolimbic dopaminergic circuits are thought to play a role in the pathophysiology of several neuropsychiatric conditions, including disorders of impulse control, schizophrenia, and neurodevelopmental disorders (Li et al., 2006; Purper-Ouakil et al., 2011; Maia and Frank, 2017; Robinson and Gradinaru, 2018), including NF1 (Brown et al., 2010; Diggs-Andrews et al., 2013; Anastasaki et al., 2015). Patients with NF1 exhibit high rates of attention deficit/hyperactivity disorder (Mautner et al., 2015; Miguel et al., 2015), in which difficulties with attentional orientation are associated with a diminished ability to suppress distractive stimuli (Aboitiz et al., 2014) such that irrelevant environmental cues are assigned exaggerated stimulus salience (Tegelbeckers et al., 2015). Previously in eLife, we showed that dopaminergic responses to light are enhanced in NF1 model mice and correlate with disruptions in the expression of conditioned behavior (Robinson et al., 2019). Our current findings suggest that these responses reflect changes in the encoding of environmental lighting conditions and, given their correlation with phenotypic expression, may reflect altered stimulus saliency. Aberrant sensory processing and motivational dysregulation are common features of neurodevelopmental disorders, including syndromic and non-syndromic forms of autism spectrum disorder (Behrmann et al., 2006; Tomchek and Dunn, 2007; Robinson and Gradinaru, 2018). Therefore, better characterization of the functional interplay between visual processing and dopaminergic circuitry may improve our pathophysiological understanding of these disorders.

## MATERIALS AND METHODS

### Key Resources Table

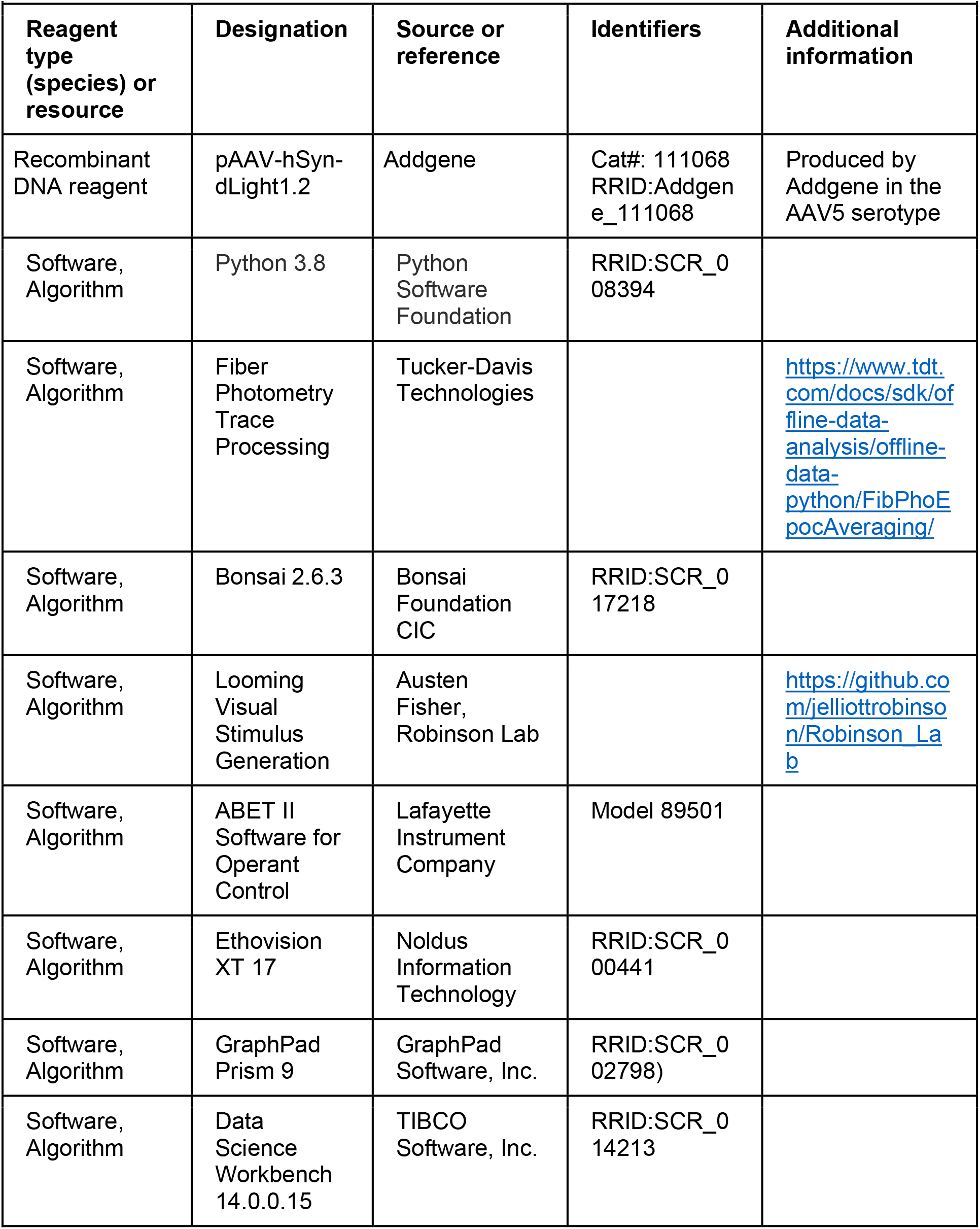
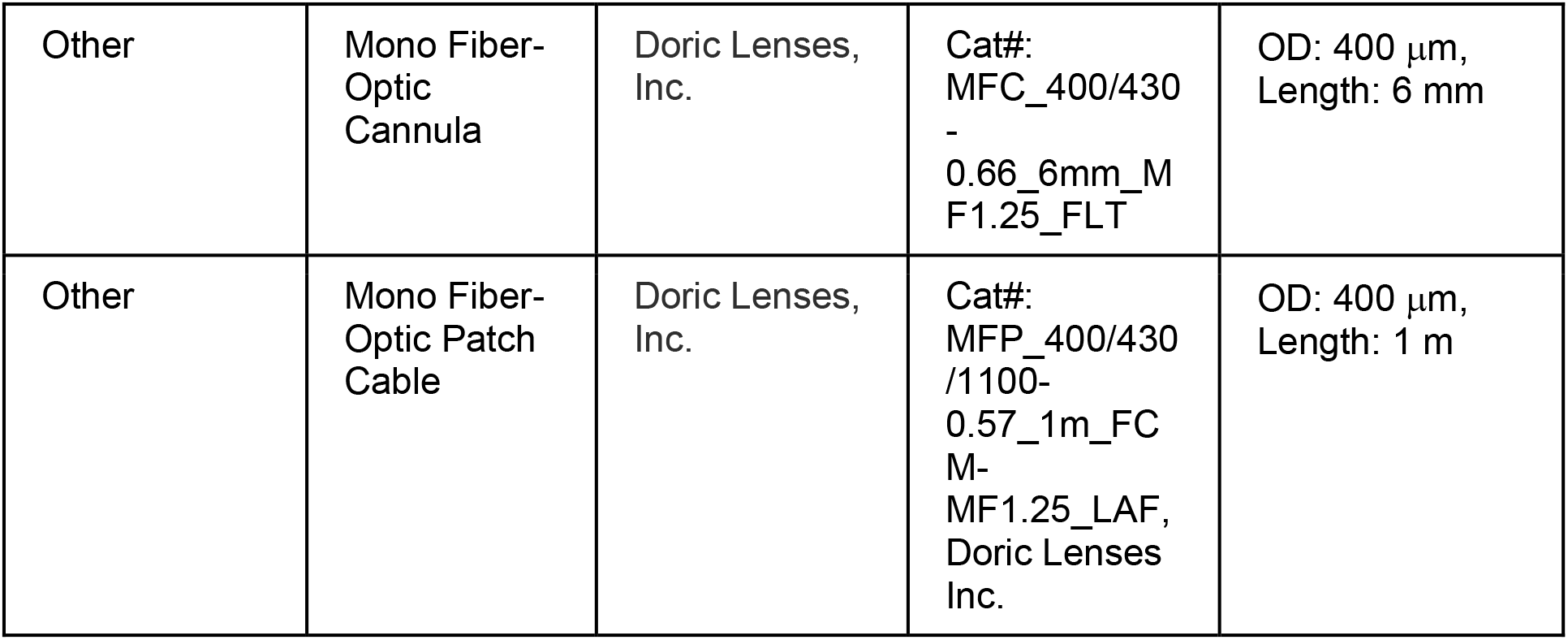

#### Experimental Animals

Experimental subjects were adult, male and female C57Bl/6J mice (the Jackson Laboratory Stock No: 000664), homozygous *Opn4* knockout mice (Panda et al., 2002), or homozygous *Gnat1/2* knockout mice (*Gnat1*^-/-^, *Gnat2*^cpfl-3^ mice; the Jackson Laboratory Stock No: 033163) that were greater than 12 weeks of age. Animals were pair or group housed (3-4 per group) throughout the duration of the experiment in a vivarium on a 14-hour/10-hour light/dark cycle (lights on at 0600 hrs, lights off at 2000 hrs) with *ad libitum* access to food and water. All experiments were performed during the light phase of the vivarium light/dark cycle, except when white LED exposure was performed 2-3 hours into the dark phase, as shown in Figure 2G. Animal husbandry and experimental procedures involving animal subjects were conducted in compliance with the Guide for the Care and Use of Laboratory Animals of the National Institutes of Health and approved by the Institutional Animal Care and Use Committee (IACUC) and by the Department of Veterinary Services at Cincinnati Children’s Hospital Medical Center (CCHMC) under IACUC protocol 2020-0058. Mice were only excluded from studies if they could not complete an entire experiment due to health concerns or loss of brain implant, if there was no dynamic photometry signal six weeks after surgery, or if the location of the photometry fiber tip was histologically determined to be outside the lateral nucleus accumbens (NAc).

#### Surgical Procedures

Stereotaxic viral vector injections and optical fiber implantation surgeries for dLight1 were performed as previously described (Robinson et al., 2019). This procedure was similar to the published protocol of Tian and colleagues (Patriarchi et al., 2019). In brief, mice were anesthetized with isoflurane (1 – 3% in 95% O_2_/5% CO_2_ provided via nose cone at 1 L/min), the scalp was shaved and sterilized with chlorhexidine surgical scrub, the skull surface was exposed, and a craniotomy hole was drilled over the lateral NAc (antero-posterior: 1.2 mm, medio-lateral: 1.6 mm relative to Bregma). 800-1000 nL of a AAV5-hSyn-dLight1.2 vector (~1 × 10^13^ viral genomes/mL, obtained from Addgene; catalog #AAV5-111068) was delivered into the LNAc (antero-posterior: 1.2 mm, medio-lateral: 1.6 mm, dorso-ventral: −4.2 mm relative to Bregma) using a blunt or beveled 34 or 35-gauge microinjection needle within a 10 uL microsyringe (NanoFil, World Precision Instruments) controlled by a microsyringe pump with SMARTouch Controller (UMP3T-1, World Precision Instruments) over ten minutes. Following viral injection, a 6 mm long, 400 μm outer diameter mono fiber-optic cannula (MFC_400/430-0.66_6mm_MF1.25_FLT, Doric Lenses Inc.) with a metal ferrule was lowered to the same stereotaxic coordinates and affixed to the skull surface with C&B Metabond (Parkel Inc.) and dental cement. Mice were given 5mg/kg carprofen (s.c.) intraoperatively and for two days postoperatively for pain. Mice were allowed a minimum of five weeks for surgical recovery and virus expression prior to participation in behavioral studies.

#### Fiber Photometry

Fluorescent signals were monitored using an RZ10x fiber photometry system from Tucker-Davis Technologies, which allowed for dLight1 excitation and emission light to be delivered and collected via the same implanted optical fiber. Our system employed a 465-nm LED for sensor excitation and a 405-nm LED for isosbestic excitation. Light was filtered and collimated using a six channel fluorescent MiniCube [FMC6_IE(400-410)_E1(460-490)_F1(500-540)_E2(555-570)_F2(580-680)_S] from Doric Lenses, Inc., which was coupled to the implanted optical fiber via a one-meter, low autofluorescence fiber optic patch cable (MFP_400/430/1100-0.57_1_FCM-MF1.25LAF, Doric Lenses Inc.). The emission signal from 405 nm isosbestic excitation was used as a reference signal to account for motion artifacts and photo-bleaching. A first order polynomial fit was applied to align the 465 nm signal to the 405 nm signal. Then, the polynomial fitted model was subtracted from the 465 nm channel to calculate ΔF values. The code for performing this function was provided by Tucker-Davis Technologies, Dr. David Root (University of Colorado, Boulder), and Dr. Marisela Morales (NIDA); it is available at: https://www.tdt.com/docs/sdk/offline-data-analysis/offline-data-python/FibPhoEpocAveraging/.

During behavioral experiments, the ΔF time-series trace was z-scored within epochs to account for data variability across animals and sessions, as described by Morales and colleagues (Barker et al., 2017). When fiber photometry was performed during sensory stimulus exposure experiments, dLight1 signals were synchronized to stimulus onset via delivery of TTL pulses to the photometry system. Peak data (magnitude, latency, and full width at half-maximal intensity) was analyzed using Python.

#### Visual Stimulus Exposure

Visual stimuli were delivered to mice during fiber photometry recordings within a custom setup that featured a 24 inch liquid crystal display (LCD) mounted 25.4 cm above mouse level in a light and sound attenuating chamber (Model 83018DDP, Lafayette Instrument Company). Stimuli (looming, static, and receding discs; full screen fades; etc.) were generated on the LCD display using Bonsai (Lopes et al., 2015), which also controlled delivery of a TTL pulse to the photometry system via a BNC cable to timestamp stimulus onset. The TTL pulse was generated with an Arduino Uno Rev3 microcontroller. During each experiment, mice were placed within the bottom of a clean shoebox cage with a thin layer cob bedding in the light and sound attenuating chamber underneath the LCD. Looming discs expanded from 0 cm to 30.5 cm over 0.84 s and froze at full expansion for 0.26 s, encompassing 61.9 degrees of visual angle, as previously described (Evans et al., 2018; Yilmaz and Meister, 2013). Receding discs shrunk from 30.5 cm to 0 cm over 0.84 s. Static discs maintained their 30.5 cm diameter throughout the duration of the stimulus. During each stimulus train, five discs were shown consecutively with a 0.5 ms interstimulus interval (ISI). Mice were exposed to five stimulus trains with a 600 s inter-trial interval (ITI). Single full field fades from black to light gray (0 – 10 s fade duration) were delivered via the LCD screen across 5 trials with a 120 s ITI.

White LED exposures were delivered via the house light of a modular conditioning chamber (Model 80015NS, Lafayette Instruments Company) placed within the light and sound attenuating box and controlled by ABET II software (Lafayette Instrument Company). A TTL breakout adapter (Model 81510) was used to synchronize stimulus delivery with the photometry recording. Single ten-second light stimuli were delivered across ten trials with a randomized ITI between 90 and 180 s. Glass neutral density filters were used to attenuate the irradiance when necessary (0.1 – 3.0 OD, HOYA Filter USA and/or Edmund Optics TECHSPEC filters). Trains of 20 one-second light stimuli with variable ISIs (10 ms – 10 s) were delivered across five trials (100 total exposures) with a 300 s ITI. One-second light stimuli with a 100 s ISI were delivered before and after 300 one-second light stimuli with a one-second ISI; 100 s separated the 300 one-second stimuli and each 100 s ISI stimulus train. One minute of 40 Hz flicker exposure (50% duty cycle) was repeated across 5 trials with a 120 s ITI. For paired light stimuli experiments, a one-second white LED stimulus was delivered one second after a 300 s or 1 s light stimulus across five trials with a 300 s ITI.

Individual wavelength light stimuli were generated with a Lumencor Aura III LED light engine, which was triggered via TTL inputs from the Lafayette Instruments TTL breakout adapter and controlled by ABET II. The liquid light guide that delivered the visual stimulus was positioned in the approximate location of the white LED within the testing chamber. LED light power (measured at mouse level with a Thor Labs PM100D optical power meter with S130VC photodiode sensor) was modulated using the onboard Lumencor graphical user interface and, when necessary, attenuated via the use of glass neutral density filters (0.1 – 3.0 OD, HOYA Filter USA and/or Edmund Optics TECHSPEC filters) placed in front of the liquid light guide outlet within a custom housing. Ten-second single wavelength stimuli were delivered in random order with a randomized ITI (140 – 200 s) to achieve five total exposures per color per mouse.

#### Auditory Stimulus Exposure

Auditory stimulus exposures were performed in the modular testing chamber within the light and sound attenuating enclosure similarly to single white LED exposures. A ten-second 80 dB tone (1 – 16 kHz; generated via Lafayette Instruments 7 Tone Generator Model 81415M) was presented via a speaker (0.25 – 16 kHz; Model 80135M14, Lafayette Instrument Company) across 5 trials with a randomized ITI (140 – 200 s).

#### Looming Stimulus Assay

The looming stimulus assay was performed as previously described (Yilmaz and Meister, 2013) using an apparatus built to the specifications of Evans et al. (2018). The apparatus featured a 20.3 cm (w) x 61 cm (l) x 40.6 cm (h) clear, open, rectangular acrylic arena with a dark, infrared (IR) light-transmitting shelter at one end and a ‘threat zone’ at the opposite end that housed a 9 cm clear plastic petri dish to encourage exploration outside of the shelter. A 15.6 inch monitor was mounted above the arena so that discs (19.5 cm maximum diameter encompassing 27 degrees of visual angle) could be presented to the mice when they entered the threat zone. The arena floor was backlit with an infrared light (880 nm back-lit collimated backlight, Advanced Illumination) to improve mouse tracking under dim light conditions. The entire apparatus was placed inside a custom light-attenuating enclosure for testing. During testing, mice were recorded with a Basler acA2040-120 um camera with a Edmunds Optics TECHSPEC 6mm C Series fixed focal length lens, and real time position tracking was performed with Bonsai. This allowed for presentation of the overhead looming, receding, or static disc stimulus to be automatically triggered when the animal was in the threat zone following a ten-minute habituation period. Mouse position and velocity data was analyzed *post hoc* using Ethovision XT software (Noldus Information Technology) and Python. Note: In Video 1, the clear, circular pedestals that separated the infrared backlight from the apparatus base can be seen with the IR camera; they were below the arena floor and inaccessible to the mouse. The setup was modified for spotlight experiments so that the pedestals would not be visible in the captured videos.

Spotlight experiments were performed in the same apparatus using the same procedure described above except that a high intensity white LED (5 mW/cm^2^ measured at mouse level) positioned to illuminate the threat zone replaced the LCD monitor.

### Statistical Analysis

Statistical analysis was performed using Python, GraphPad Prism 9 (GraphPad Software, Inc.), and/or Data Science Workbench 14 (for 3-way repeated measures ANOVA; TIBCO Software, Inc.). All statistical tests performed on data presented in the manuscript are stated in the figure captions and provided in detail with the corresponding source data in the Supplemental Data and Statistical Analysis file. For each experiment, statistical tests were chosen based on the structure of the experiment and data set. No outliers were removed during statistical analysis. Parametric tests were used throughout the manuscript. Sample size estimates were based on studies in Robinson et al. (2019) and power analysis performed using the sampsizepwr() function in Matlab (MathWorks). When analysis of variance (ANOVA; 1-way, 2-way, 3-way, and/or repeated measures) was performed, multiple comparisons were corrected using the Bonferroni correction. When repeated measures ANOVA could not be performed due to missing values (Figure 2-Figure Supplement 1, panel C), data was analyzed by fitting a mixed model in GraphPad Prism 9; this approach uses a compound symmetry covariance matrix and is fit using restricted maximum likelihood (REML). When results were compared to a pre-stimulus baseline, this value was defined as the amplitude of the dLight1 peak that occurred 500 ms prior to stimulus delivery. When results were compared to a ‘null’ stimulus, the value was defined as the dLight1 peak that occurred at the onset of a TTL that timestamped a trial in which no stimulus was delivered.

## Data and Materials Availability

Viral vector plasmids used in this study are available on Addgene. Codes used for fiber photometry signal extraction and analysis are available at https://www.tdt.com/docs/sdk/offline-data-analysis/offline-data-python/FibPhoEpocAveraging/. Codes used for visual stimulus generation are available at https://github.com/jelliottrobinson/Robinson_Lab. Source data is available in the provided Supplemental Data and Statistical Analysis file.

## ACKNOWLEDGEMENTS

We would like to acknowledge Mary Claire Casper and Shiva Senthilkumar for assistance with histological sample preparation and preliminary data analysis, respectively. We would also like to thank Dr. Gregory Schwartz at Northwestern University Feinberg School of Medicine and Dr. Diego Fernandez at the National Institute of Mental Health for helpful discussions regarding technical considerations and/or interpretation of the experimental findings. This work was funded by a Cincinnati Children’s Research Foundation Trustee Award, a Simons Foundation Autism Research Initiative (SFARI) Bridge to Independence Award (663007), a SFARI Supplement to Enhance Equity and Diversity (SEED) Award, and a Gilbert Family Foundation Neurofibromatosis Gene Therapy Initiative Team Science Award to JER.

## VIDEO TITLES

**Video 1. Behavioral response to presentation of black looming discs on a light background when mice entered the threat zone of a rectangular arena.**

**Video 2. Behavioral response to illumination of a spotlight when mice entered the target zone of a rectangular arena.**

---

**Figure 2-Figure Supplement 1.**
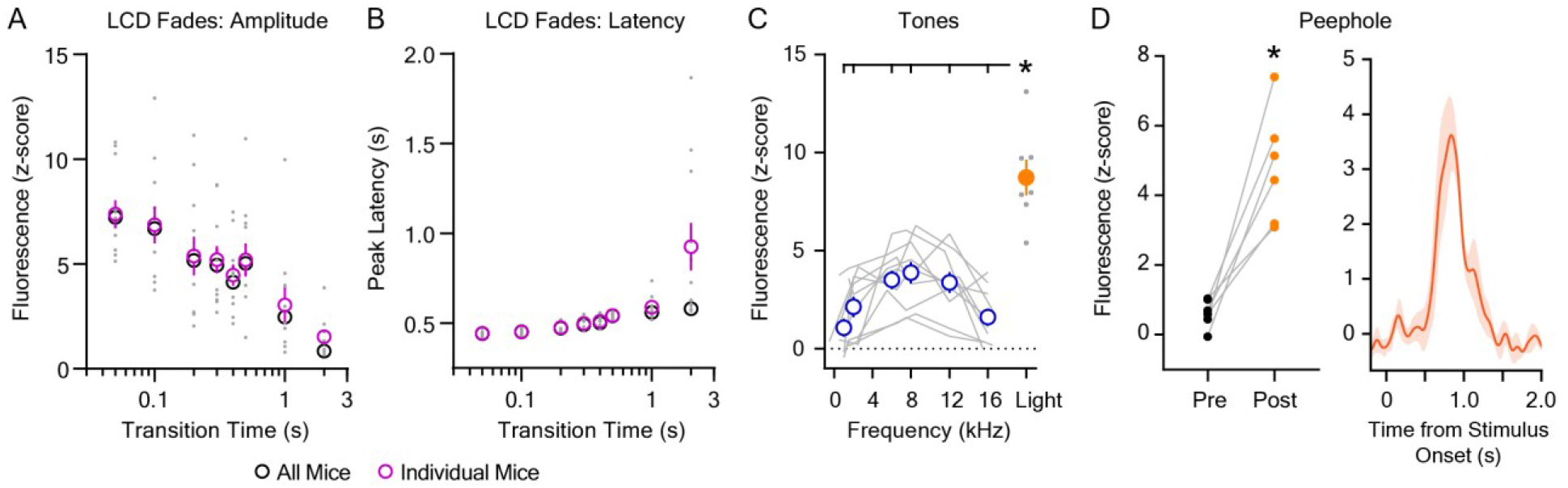
**A.** The magnitude of the dopaminergic response to dark-to-light transitions was dependent on the stimulus transition time (n = 11; 1-way repeated measures ANOVA; F_7,70_ = 17.76, p < 0.001). **B.** The latency of the peak dopaminergic response to dark-to-light transitions was dependent on the stimulus transition time (n = 11; 1-way repeated measures ANOVA; F_7,70_ = 11.03, p = 0.007). In A and B, pink circles (mean ± SEM) and gray dots (individual values) represent data derived from fluorescent dLight1 traces averaged across trials for each individual mice. Shown for comparison, black circles represent data derived from fluorescence traces averaged across all mice prior to peak detection, which yielded a single value for each transition time. **C.** The dopaminergic response to 80 dB tones was dependent on their frequency (n = 10; 1-way repeated measures ANOVA; F_5,45_ = 10.11, p < 0.001) and was less than the response to a 5 μW/cm^2^ LED light stimulus (Mixed-effects ANOVA; F = 26.62, p_stimulus_ < 0.001). Bonferroni *post hoc* tests revealed that the dLight1 response to the light stimulus was significantly larger than all tone responses. **D.** (*Left*) Opening the enclosure peephole evoked significant dopamine release at stimulus onset when compared to the pre-stimulus baseline (n = 6; paired t-test; t5 = 5.72, p = 0.002). (*Right*) Averaged dLight1 trace showing LNAc dopamine evoked by opening the peephole on the door of the behavioral testing chamber enclosure to observe mouse behavior ± SEM. The irradiance associated with this manipulation was 17 nW/cm^2^ measured at mouse level. In all panels, * indicates p < 0.05.

**Figure 4-Figure Supplement 1.**
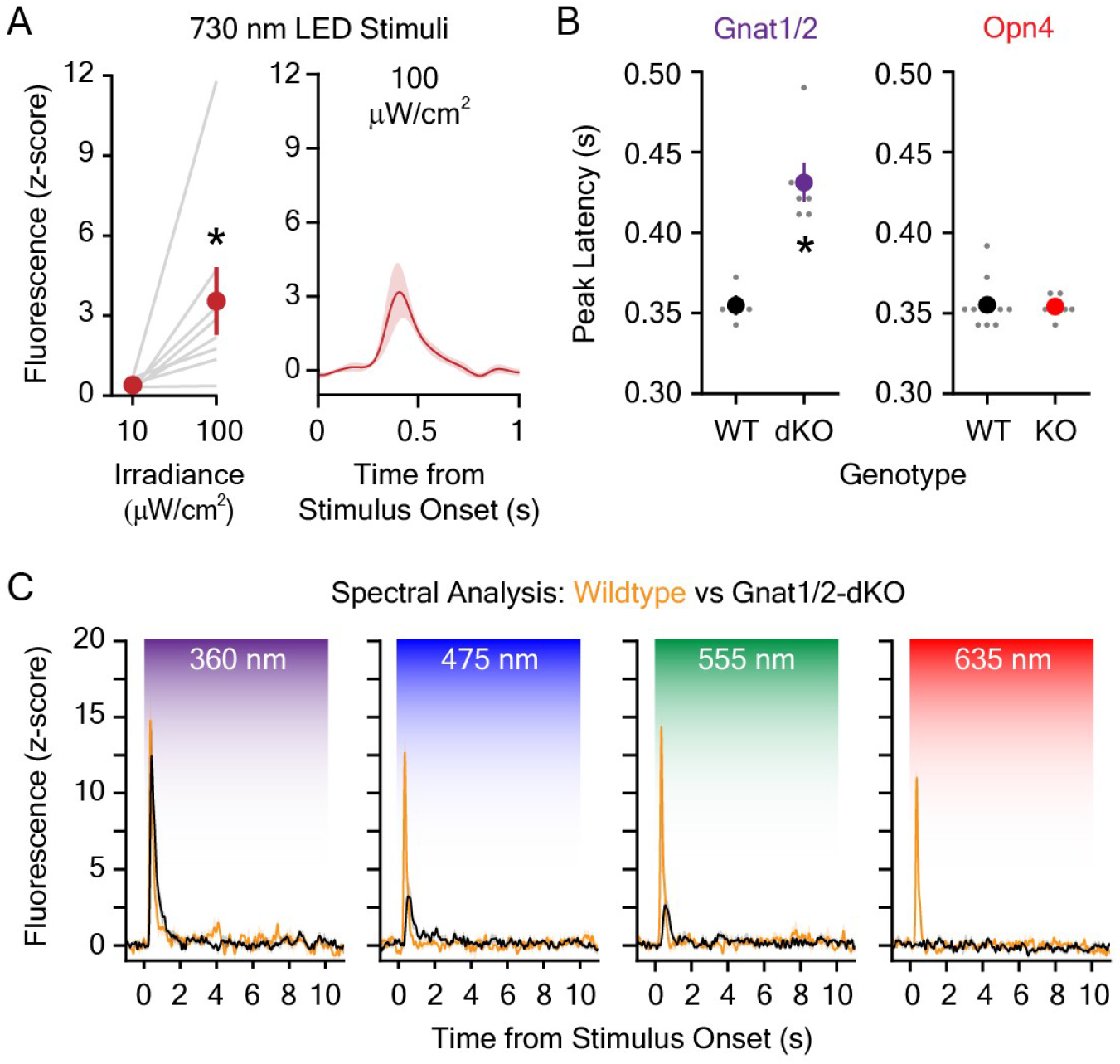
**A.** (*Left*) 100 μW/cm^2^ far-red (730 nm) LED light evoked significantly greater dopamine release compared to 10 μW/cm^2^ light (n = 8; paired t-test; t7 = 2.54, p = 0.04). (*Right*) Averaged dLight1 trace showing LNAc dopamine evoked by a 100 μW/cm^2^ far-red LED ± SEM. **B.** (*Left*) The latency of the peak dLight1 response to 5.0 μW/cm^2^ white light was significantly greater in *Gnat1/2* double knockout (dKO) mice relative to wildtype controls (WT; n_WT_ = 4, n_KO_ = 6; unpaired t-test; t8 = 4.77, p = 0.001). (*Right*) There was no difference in the latency of the peak dLight1 response to 5.0 μW/cm^2^ white light in *Opn4* knockout mice (KO) relative to wildtype controls (WT; *black;* n_WT_ = 11, n_KO_ = 6; unpaired t-test; t_15_ = 0.16, p = 0.87). **C.** Averaged dLight1 trace showing LNAc dopamine evoked by 10 μW/cm^2^ UV (360 nm), blue (475 nm), green (555 nm), or red (635) LEDs ± SEM in wildtype (orange) or *Gnat1/2-dKO* (black) mice. * indicates p < 0.05.

